# Bidirectional Long-Term Synaptic Zinc Plasticity at Mouse Glutamatergic Synapses

**DOI:** 10.1101/320671

**Authors:** Nathan W. Vogler, Thanos Tzounopoulos

## Abstract

Synaptic zinc is coreleased with glutamate to modulate neurotransmission in many excitatory synapses. In the auditory cortex, synaptic zinc modulates sound frequency tuning and enhances frequency discrimination acuity. In auditory, visual, and somatosensory circuits, sensory experience causes long-term changes in synaptic zinc levels and/or signaling, termed here *synaptic zinc plasticity*. However, the mechanisms underlying synaptic zinc plasticity and the effects of this plasticity on long-term glutamatergic plasticity remain unknown. To study these mechanisms, we used male and female mice and employed *in vitro* and *in vivo* models in zinc-rich, glutamatergic dorsal cochlear nucleus (DCN) parallel fiber (PF) synapses. High-frequency stimulation of DCN PF synapses induced long-term depression of synaptic zinc signaling (Z-LTD), as evidenced by reduced zinc-mediated inhibition of AMPA receptor (AMPAR) excitatory postsynaptic currents (EPSCs). Low-frequency stimulation induced long-term potentiation of synaptic zinc signaling (Z-LTP), as evidenced by enhanced zinc-mediated inhibition of AMPAR EPSCs. Thus, Z-LTD is a new mechanism of LTP and Z-LTD is a new mechanism of LTP. Pharmacological inhibition of Group 1 metabotropic glutamate receptors (G1 mGluRs) eliminated Z-LTD and Z-LTP. Pharmacological activation of G1 mGluRs induced Z-LTD and Z-LTP, associated with bidirectional changes in presynaptic zinc levels. Finally, exposure of mice to loud sound caused G1 mGluR-dependent Z-LTD in DCN PF synapses, consistent with our *in vitro* results. Together, we show that G1 mGluR activation is necessary and sufficient for inducing bidirectional long-term synaptic zinc plasticity.

**Key points summary:**

- Synaptic zinc is coreleased with glutamate to modulate neurotransmission and auditory processing. Sensory experience causes long-term changes in synaptic zinc signaling, termed synaptic zinc plasticity.
- At zinc-containing glutamatergic synapses in the dorsal cochlear nucleus (DCN), we show that high-frequency stimulation reduces synaptic zinc signaling (Z-LTD), whereas low-frequency stimulation increases synaptic zinc signaling (Z-LTP).
- Group 1 metabotropic glutamate receptor (mGluR) activation is necessary and sufficient to induce Z-LTP and Z-LTD. Z-LTP and Z-LTD are associated with bidirectional changes in presynaptic zinc levels.
- Sound-induced Z-LTD at DCN synapses requires Group 1 mGluR activation.
- Bidirectional synaptic zinc plasticity is a previously unknown mechanism of LTP and LTD at zinc-containing glutamatergic synapses.

## Introduction

In many brain areas, including the neocortex, limbic structures, and the auditory brainstem, glutamatergic vesicles are loaded with zinc (Danscher & Stoltenberg, 2005; Frederickson *et al.*, 2005). This pool of mobile, synaptic zinc is coreleased with glutamate. Synaptically released zinc inhibits synaptic and extrasynaptic NMDA receptor (NMDAR) EPSCs, and modulates AMPA receptor (AMPAR) EPSCs (Vogt *et al.*, 2000; Vergnano *et al.*, 2014; Anderson *et al.*, 2015; Kalappa *et al.*, 2015; Kalappa & Tzounopoulos, 2017). Namely, synaptic zinc inhibits AMPAR EPSCs during baseline synaptic activity via postsynaptic mechanisms, but enhances steady-state AMPAR EPSCs during higher frequencies of synaptic stimulation (Kalappa *et al.*, 2015; Kalappa & Tzounopoulos, 2017). The enhancing effect of synaptic zinc on AMPAR EPSCs is short-lasting and is mediated by short-term, zinc-mediated changes in presynaptic glutamatergic neurotransmission (Perez-Rosello *et al.*, 2013; Kalappa & Tzounopoulos, 2017). Thus, synaptic zinc is a major modulator of baseline neurotransmission and short-term plasticity of glutamatergic synapses.

In awake mice, synaptic zinc enhances the responsiveness (gain) of auditory cortical principal neurons to sound, but reduces the gain of cortical interneurons (Anderson *et al.*, 2017). Furthermore, synaptic zinc sharpens the sound frequency tuning of auditory cortical principal neurons, and enhances frequency discrimination acuity (Kumar *et al.*, 2019). Sensory experience bidirectionally modulates the levels of vesicular zinc and synaptic zinc signaling in several sensory brain areas (Nakashima & Dyck, 2009; Kalappa *et al.*, 2015; Li *et al.*, 2017; McAllister & Dyck, 2017). In the somatosensory cortex, whisker plucking increases zinc levels, whereas whisker stimulation reduces zinc levels (Brown & Dyck, 2002, 2005). In the primary visual cortex, monocular deprivation increases vesicular zinc levels (Dyck *et al.*, 2003). In the retina, optic nerve damage increases zinc levels, which in turn inhibit optic nerve regeneration and promote cell death (Li *et al.*, 2017). In the dorsal cochlear nucleus (DCN), an auditory brainstem nucleus, exposure to loud sound reduces vesicular zinc levels and synaptic zinc signaling (Kalappa *et al.*, 2015). Yet, the cellular and molecular mechanisms underlying the long-term experience-dependent plasticity of synaptic zinc signaling, termed here *synaptic zinc plasticity*, and the relationship of synaptic zinc plasticity to long-term glutamatergic synaptic plasticity remain unknown. Elucidating these mechanisms is crucial for understanding how the brain adapts during normal sensory processing, and why it fails to properly adjust in sensory disorders associated with pathological central adaptation, such as in tinnitus (Auerbach *et al.*, 2014).

To determine the mechanisms of long-term synaptic zinc plasticity and its effects on LTP and LTD, we developed *in vitro* and *in vivo* models. Namely, we used electrophysiology, pharmacology, and fluorescent imaging in the DCN, which contains granule cell endings, parallel fibers (PFs), with high levels of synaptic zinc (Frederickson *et al.*, 1988; Rubio & Juiz, 1998; Kalappa *et al.*, 2015). We investigated these mechanisms *in vitro*, in response to electrical synaptic activation that induces synaptic plasticity such as LTP and LTD in brain slices, as well as *in vivo*, in response to loud sound exposure. Our results demonstrate that bidirectional activity-dependent synaptic zinc plasticity is a previously unknown, Group 1 mGluR-dependent mechanism of LTP and LTD at zinc-containing glutamatergic synapses.

## Materials and Methods

### Animals

Male or female ICR mice (Envigo) were used in this study, aged between postnatal day 17 (P17) to P28. All animal procedures were approved by the Institutional Animal Care and Use Committee of the University of Pittsburgh, Pittsburgh, PA.

### Brain slice preparation

Mice were deeply anesthetized with isoflurane (3% in O_2_), then immediately decapitated and their brains were removed. Brain slices were prepared in artificial cerebrospinal fluid (ACSF, 34°C) containing the following (in mM): 130 NaCl, 3 KCl, 1.2 CaCl_2_ 2H_2_O, 1.3 MgCl_2_6H_2_O, 20 NaHCO_3_, 3 HEPES, and 10 D-Glucose, saturated with 95% O_2_/5% CO_2_ (vol/vol), pH = 7.25-7.35, ∼300 mOsm. Using a Vibratome (VT1200S; Leica), coronal brain slices (210 μm thickness) containing the left dorsal cochlear nucleus (DCN) were cut, then placed in a chamber containing warm (34°C) ACSF, and incubated for 60 min at 34°C, then room temperature (no longer than 3 hours) before beginning electrophysiology experiments. Incubating ACSF was the same as cutting ACSF, except it was stirred with Chelex 100 resin (Bio-Rad) for 1 hour to remove contaminating zinc, then filtered using Nalgene rapid flow filters lined with polyethersulfone (0.2 μm pore size). After filtering, high purity CaCl_2_ 2H_2_O and MgCl_2_6H_2_O (99.995%; Sigma Aldrich) were added. All plastic and glassware used for these experiments were washed with 5% nitric acid.

### Electrophysiology

*Whole-cell recordings*. DCN slices were transferred to the recording chamber and perfused with ACSF (1-2 mL/min), maintained at ∼34°C using an inline heating system (Warner Instruments). Recording ACSF was the same as incubating ACSF (see above), except it contained 2.4 mM CaCl_2_ 2H_2_O. Whole-cell recordings from cartwheel cells were performed using glass micropipettes (3-6 MΩ; Sutter Instruments). Cartwheel cells were identified by the presence of complex spikes in cell-attached configuration before break-in or in response to current injections in current-clamp mode after break-in (Zhang & Oertel, 1993; Manis *et al.*, 1994; Tzounopoulos *et al.*, 2004). Recording pipettes were filled with a potassium-based internal solution (except for Figure 6, see below) containing the following (in mM): 113 K-gluconate, 4.5 MgCl_2_6H_2_O, 14 Tris-phosphocreatine, 9 HEPES, 0.1 EGTA, 4 Na_2_ATP, 0.3 Tris-GTP, and 10 sucrose (pH = 7.25, 295 mOsm). For experiments shown in Figure 6 measuring NMDAR EPSCs, recordings were performed using a cesium-based internal solution containing the following (in mM): 128 Cs(CH_3_O_3_S), 10 HEPES, 4 MgCl_2_6H_2_O, 4 Na_2_ATP, 0.3 Tris-GTP, 10 Tris-phosphocreatine, 1 EGTA, 1 QX-314, and 3 Na-ascorbate (pH = 7.25, 300 mOsm). Voltages were not corrected for junction potentials. Recordings were performed using ephus (Suter *et al.*, 2010) and a MultiClamp 700B amplifier (Axon Instruments). Data were sampled at 10 kHz and low-pass-filtered at 4 kHz. Series resistance (R_s_) and input resistance (R_m_) were monitored during the recording period by delivering −5 mV voltage steps for 50 ms. R_s_ was calculated by dividing the −5 mV voltage step by the peak current generated immediately after the voltage step. R_m_ was calculated by dividing the −5 mV voltage step by the difference between the baseline and steady-state hyperpolarized current, then subtracting R_s_. Data were excluded if R_s_ or R_m_ changed by more than 20% from the baseline period. EPSCs were evoked using an Isoflex stimulator (A.M.P.I., 0.1 ms pulses) through a glass ACSF-containing theta electrode to stimulate the zinc-rich parallel fibers. All EPSCs were recorded in the presence of SR95531 (20 μM, GABA_A_R antagonist) and strychnine (1 μM, GlyR antagonist). AMPAR EPSCs were recorded in voltage-clamp mode at −70 mV. For paired-pulse experiments, the inter-stimulus interval was 50 ms. NMDAR EPSCs were evoked by a 5-pulse stimulus train (20 Hz) (Anderson *et al.*, 2015), recorded in voltage clamp mode at +40 mV, and in the presence of DNQX (20 μM, AMPA/kainate receptor antagonist). All drugs were always bath applied.

*Induction of plasticity*. High-frequency stimulation (HFS) consisted of 3 trains of 100 Hz pulses for 1 sec, with 10 sec between trains. For the experiments shown in Figure 1 C, a subset of cells (n=5) were depolarized to −10 mV during each HFS train, while the other subset (n=6) were held at −70 mV during HFS. Because we observed no difference in the zinc plasticity (% potentiation by ZX1) between these subsets (depolarized = 2.68 ± 5.36%, non-depolarized = 8.76 ± 8.64%, *p* = 0.58, unpaired *t* test), they were grouped together for subsequent analysis. ZX1 (100 μM) is a fast, high-affinity extracellular zinc chelator (Anderson *et al.*, 2015; Kalappa *et al.*, 2015; Kalappa & Tzounopoulos, 2017). For all other experiments, cells were voltage-clamped at −70 mV during HFS. For experiments measuring NMDAR EPSCs after HFS (Figure 6), DNQX (20 μM) was added after HFS, then cells were voltage-clamped at +40 mV to record NMDAR EPSCs. For ifenprodil experiments (Figure 6 D-E), ZX1 was applied prior to ifenprodil to chelate extracellular zinc, because zinc affects NMDAR ifenprodil sensitivity (Hansen *et al.*, 2014). In these experiments after HFS, ZX1 was applied with DNQX, after the HFS. Low-frequency stimulation (LFS) consisted of 5 Hz pulses for 3 min. During LFS, cells were held at −80 mV in current-clamp mode. To isolate mGluR-mediated plasticity, all LFS experiments were performed in the presence of APV (50 μM, NMDAR antagonist), and with external ACSF containing 4 mM CaCl_2_ 2H_2_O and 4 mM MgCl_2_6H_2_O (Oliet *et al.*, 1997). The interleaved experiments shown in Figure 4 D, examining the effect of 50 μM DHPG application, were also performed in these conditions. For normalized EPSCs (% baseline), EPSC amplitudes were normalized to the average EPSC amplitude during the 5 min baseline period before HFS/LFS, DHPG, ifenprodil, or ZX1 application. To quantify ZX1 potentiation after HFS/LFS or DHPG application, EPSC amplitudes were renormalized to the average EPSC amplitude of the new baseline period 5 min before ZX1 application. ZX1 potentiation (shown in bar graphs) was quantified as the percent increase in the average EPSC amplitude during the last 5 min of ZX1 application compared to the 5 min baseline period before ZX1 application.

**Figure 1.**
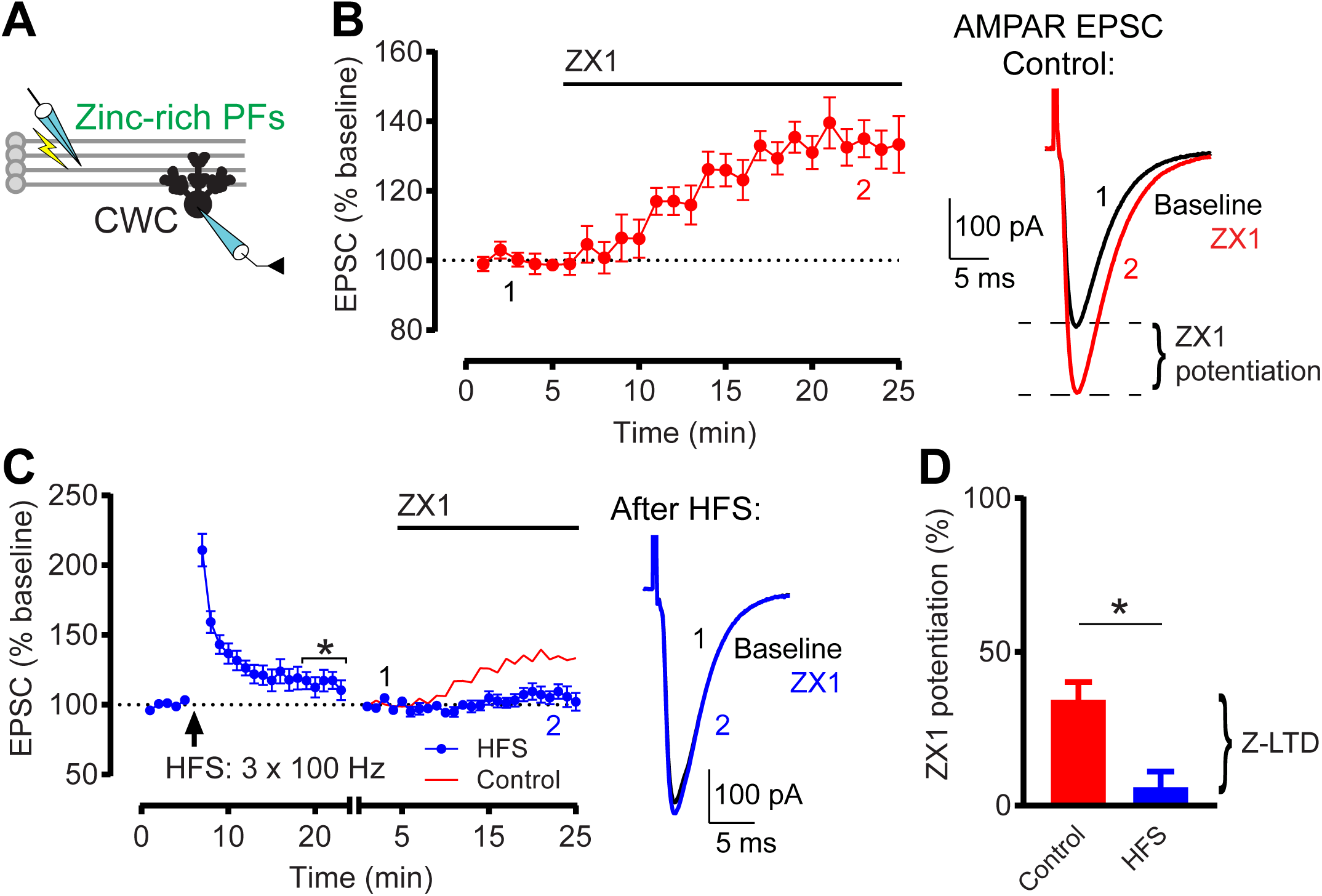
High-frequency stimulation (HFS) induces Z-LTD in DCN parallel fiber synapses. **(A)** Schematic of experimental setup illustrating stimulation of zinc-rich glutamatergic DCN parallel fibers (PFs) and whole-cell recording of a postsynaptic cartwheel cell (CWC). **(B)** *Left:* Time course of AMPAR EPSC amplitude before and after ZX1 application, normalized to baseline before ZX1 application (100 μM). *Right:* Example AMPAR EPSCs before and after ZX1 application, showing ZX1 potentiation. **(C)** *Left:* Time course of AMPAR EPSC amplitude before and after HFS, and before and after subsequent ZX1 application (blue). After obtaining a stable baseline, HFS was delivered (3 × 100 Hz for 1 sec, 10 sec ISI). EPSC % baseline after HFS (mins. 19-23): n=11, *p=0.041, one-sample t test vs. 100%. Star (*) indicates significant LTP. To examine ZX1 potentiation after HFS, after obtaining a stable baseline after HFS, AMPAR EPSC amplitude was renormalized to the new baseline before ZX1 application. The renormalization is indicated by a gap and restart of timing in the x-axis. For comparison, red line shows normalized time course of AMPAR EPSC amplitude before and after ZX1 application in controls replotted from **B**. *Right:* Example AMPAR EPSCs showing no ZX1 potentiation after HFS. **(D)** Average ZX1 potentiation (% increase from baseline) during the last 5 min of ZX1 application. ‘Control’ (n=10) vs. ‘HFS’ (n=11): *p=0.002, unpaired t test. The reduction in ZX1 potentiation is termed Z-LTD. Values represent mean ± SEM. Star (*) indicates p<0.05. For detailed values and statistical tests for all figures, see Materials and Methods, Statistical Analysis section.

### Vesicular zinc imaging with DA-ZP1

After preparation and incubation of DCN slices (described above), slices were transferred to the imaging chamber and perfused with recirculating ACSF (2-3 mL/min) maintained at ∼34°C. Imaging of presynaptic vesicular zinc levels in DCN parallel fibers was performed using DA-ZP1, a high-affinity, membrane permeable fluorescent zinc sensor (Zastrow *et al.*, 2016). DA-ZP1 (0.5-1.0 µM) was added to the ACSF, and allowed to incubate for at least 20 min before imaging. Images were acquired using an upright microscope (Olympus BX5) with epifluorescence optics through a 20x water immersion objective (Olympus). Green fluorescent signals were isolated using a Pinkel filter set (Semrock LF488/543/625-3X-A-000) in response to excitation by an ephus-driven blue LED (M470L2; Thorlabs), and images were acquired using a CCD camera (Retiga 2000R, QImaging). Images consisted of 20 frames captured at 0.067 Hz which were then averaged together and analyzed in MATLAB (Mathworks). The DCN molecular layer, which contains the vesicular zinc-rich parallel fibers, extends ∼75 µm deep from the ependymal surface, while deeper layers lack vesicular zinc (zinc-free region) (Ryugo & Willard, 1985; Frederickson *et al.*, 1988; Rubio & Juiz, 1998). Thus, DA-ZP1 produces a band of fluorescence within the molecular layer near the ependymal surface, consistent with the distribution of zinc-rich parallel fiber terminals (Frederickson *et al.*, 1988; Kalappa *et al.*, 2015; Zastrow *et al.*, 2016). The DA-ZP1 fluorescence band is absent in ZnT3 KO mice lacking vesicular zinc, indicating that it specifically labels vesicular zinc (Kalappa *et al.*, 2015; Zastrow *et al.*, 2016). To control for slice-to-slice variability in the molecular layer volume, which in turn might lead to variability in DA-ZP1 brightness, we compared DA-ZP1 fluorescence in the same region of the same slice before and after DHPG application (Figure 5). DA-ZP1 fluorescence 15-20 min after DHPG application was normalized to baseline fluorescence before DHPG application. To quantify DA-ZP1 fluorescence, we quantified two ROIs within each slice: one within the zinc-containing molecular layer (zinc ROI) and the other within the zinc-free region (zinc-free ROI) (Kalappa *et al.*, 2015; Zastrow *et al.*, 2016). Because the DCN molecular layer is curved along the ependymal surface, to define the zinc ROI, we used a MATLAB routine to automatically detect the abrupt increase in fluorescence intensity between the background and the ependymal surface of the slice. Then the zinc ROI was automatically selected to include 50 µm depth from the ependymal surface, consistent with the extent of the zinc-containing parallel fiber terminals (Frederickson *et al.*, 1988). The length of the ROI was 450 µm. The zinc-free ROI was identical to the zinc ROI, except located 200-250 µm from the border of the slice, within the zinc-free region (deep or fusiform cell layers) (Ryugo & Willard, 1985; Frederickson *et al.*, 1988). Thus, all ROIs contained the same cross-sectional area. The automatically generated ROI borders are shown with yellow lines in Figure 5. Fluorescence intensity was averaged within each ROI, and the zinc-sensitive fluorescence was calculated by subtracting the zinc-free ROI fluorescence from the zinc ROI fluorescence.

### Noise exposure

Noise exposure was performed based on previously published methods (Kalappa *et al.*, 2015). Sham-or noise-exposed mice were anesthetized using 3% isoflurane during induction and 1-1.5% during maintenance. Noise-exposed mice were exposed for 4 hours to narrow bandpass noise at 116 dB sound pressure level (SPL), centered at 16 kHz with a 1.6 kHz bandwidth. Noise was presented unilaterally (left ear) through a pipette tip inserted into the left ear canal, with the other end attached to a calibrated speaker (CF-1; Tucker Davis Technologies). Insertion of the pipette tip into the ear canal did not produce a seal. Sham-exposed mice underwent an identical procedure except without any noise exposure. For mice given intraperitoneal injections of AIDA (2 mg/kg), one injection was given 30 min prior to exposure, and a second injection was given 2 hours later. After noise-or sham-exposure, ABRs were collected and mice recovered from anesthesia, then DCN slices were prepared (within 30 min after exposure).

### ABRs

Auditory Brainstem Responses (ABRs) were measured based on previously published methods (Kalappa *et al.*, 2015). ABRs were recorded immediately after noise-or sham-exposure. During ABR measurements, mice were anesthetized using 3% isoflurane during induction and 1-1.5% during maintenance. Mice were placed in a sound attenuating chamber and temperature was maintained at ∼37°C using a heating pad. A subdermal electrode was placed at the vertex, the ground electrode placed ventral to the right pinna, and the reference electrode placed ventral to the left pinna (sham- or noise-exposed ear). In noise-exposed mice, because no ABRs were detected when recording from the exposed (ipsilateral) ear, we recorded ABRs from the non-exposed (contralateral) ear (Figure 7 D). For ABR measurements from contralateral ears of noise-exposed mice, the reference electrode was placed ventral to the right pinna (contralateral ear) and the ground electrode placed ventral to the left pinna. ABRs were detected in response to 1 ms click sound stimuli, presented through a pipette tip inserted into the ear canal, with the other end attached to the speaker (CF-1; Tucker Davis Technologies). ABRs were recorded in response to clicks presented in 10 dB steps, ranging from 0-80 dB SPL. 1 ms clicks were presented at a rate of 18.56/sec using System 3 software package from Tucker Davis Technologies, and ABRs were averaged 512 times and filtered using a 300-3,000 Hz bandpass filter. ABR threshold was defined as the lowest stimulus intensity which generated a reliable Wave 1 in the response waveform. Wave 1 amplitude was measured as the peak-to-trough amplitude of the first wave in the ABR waveform (latency ∼2 ms), in response to 80 dB SPL clicks.

### Drugs

All chemicals used for ACSF and internal solutions were purchased from Sigma-Aldrich. The following drugs were purchased from HelloBio: SR95531 hydrobromide, DL-AP5, DNQX disodium salt, ifenprodil, MPEP hydrochloride, LY367385, and (S)-3,5-Dihydroxyphenylglycine (DHPG). Strychnine hydrochloride was purchased from Abcam. (RS)-1-Aminoindan-1,5-dicarboxylic acid (AIDA) was purchased from Tocris. ZX1 was purchased from STREM Chemicals.

### Statistical Analysis

All data analysis was performed using Matlab (Mathworks), Excel (Microsoft), or Prism 7 (GraphPad). For statistical tests for normalized data, or within groups, we used one-sample *t* tests (for normally distributed data) or Wilcoxon signed rank tests (for non-normally distributed data). Data were considered normally distributed if they passed the Shapiro-Wilk normality test. For comparisons between two (normally distributed) groups, we used unpaired *t* tests. All *t* tests were two-tailed. For comparisons between three groups, we used ordinary one-way ANOVA with Bonferroni’s multiple comparisons test (for normally distributed data), or Kruskal-Wallis test with Dunn’s multiple comparisons test (for non-normally distributed data). IC_50_ was calculated using the Hill equation by fitting the dose-response curve with a nonlinear least squares fit. The IC_50_ of each fit was compared using the extra sum-of-squares F test. Significance levels are defined as *p* < 0.05. Group data are presented as mean ± SEM.

*Detailed values and statistical tests for Figures*. Figure 1: **(1C)** EPSC % baseline after HFS (average of mins. 19-23): 115.1 ± 6.43%, n=11, t=2.34 df=10, *p=0.041, one-sample t test vs. 100%. **(1D)** ZX1 potentiation (%): ‘Control’: 34.47 ± 5.7%, n=10, t=6.049 df=9, *p=0.0002, one-sample t test vs. 0%. ‘HFS’: 6.0 ± 5.15%, n=11, t=1.165 df=10, n.s. p=0.27, one-sample t test vs. 0%. ‘Control’ vs. ‘HFS’: t=3.719 df=19, *p=0.0015, unpaired t test.

Figure 2: **(2A)** EPSC % baseline after HFS (average of mins. 19-23): 124.8 ± 4.76%, n=9, t=5.198 df=8, *p=0.0008, one-sample t test vs. 100%. **(2B)** EPSC % baseline after HFS (average of mins. 19-23): 126.2 ± 9.54%, n=6, t=2.743 df=5, *p=0.041, one-sample t test vs. 100%. One cell was included for analysis of EPSCs following HFS, but did not remain stable throughout subsequent ZX1 application and was excluded from analysis following ZX1 application, quantified in C. **(2C)** ZX1 potentiation (%): ‘HFS + APV’: 4.28 ± 6.08%, n=9, t=0.7039 df=8, n.s. p=0.502, one-sample t test vs. 0%. ‘HFS + LY367385, MPEP, APV’: 36.07 ± 9.05%, n=5, t=3.987 df=4, *p=0.016, one-sample t test vs. 0%. One-way ANOVA: F= 7.737, *p=0.003. ‘Control’ vs. ‘HFS + APV’: *p=0.0038; ‘HFS + APV’ vs. ‘HFS + LY367385, MPEP, APV’: *p= 0.0115; Bonferroni’s multiple comparisons test.

**Figure 2.**
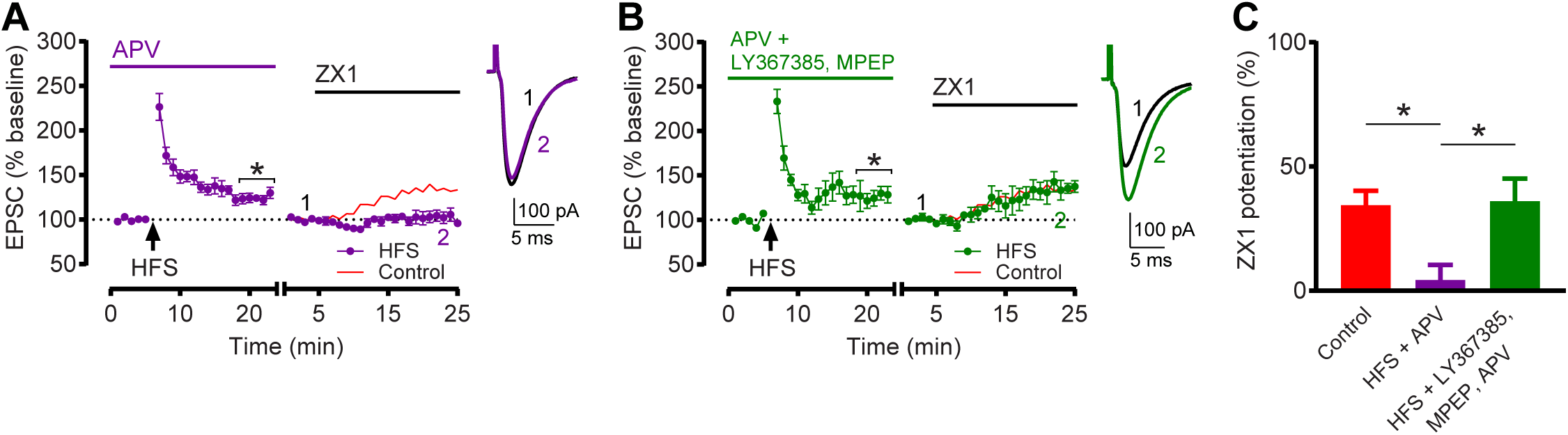
Group 1 mGluR activation is required for HFS-induced Z-LTD. **(A)** Time course of AMPAR EPSC amplitude before and after HFS in the presence of APV (50 μM), and before and after subsequent ZX1 application. EPSC % baseline after HFS (mins. 19-23): n=9, *p=0.0008, one-sample t test vs. 100%. **(B)** Same time course as in **A** but in the presence of LY367385 (100 μM), MPEP (4 μM), and APV (50 μM). EPSC % baseline after HFS (mins. 19-23): n=6, *p=0.041, one-sample t test vs. 100%. For **(A-B)**, star (*) indicates significant LTP. To examine ZX1 potentiation after HFS, similar approach and renormalization as in **1C** was performed. Red line shows the time course of AMPAR EPSC amplitude before and after ZX1 application in control replotted from **1B**. Example traces show AMPAR EPSCs before and after ZX1. **(C)** Average ZX1 potentiation (% increase from baseline) during the last 5 min of ZX1 application, with control data from **1D.** ‘HFS + APV’ (n=9) reduced ZX1 potentiation compared to control; this reduction was blocked by LY367385 and MPEP (n=5). One-way ANOVA/Bonferroni, *p=0.003. Values represent mean ± SEM. Star (*) indicates p<0.05.

Figure 3: **(3A)** EPSC % baseline after LFS (average of mins. 19-23): 95.97 ± 3.5%, n=8, t=1.155 df=7, n.s. p=0.29, one-sample t test vs. 100%. Two cells were included for analysis of EPSCs following LFS, but did not remain stable throughout subsequent ZX1 application and were excluded from analysis following ZX1 application, quantified in C. **(3B)** EPSC % baseline after LFS (average of mins. 20-24): 73.67 ± 5.8%, n=6, t=4.528 df=5, *p=0.006, one-sample t test vs. 100%. **(3C)** ZX1 potentiation (%): ‘Control’: 19.65 ± 4.3%, n=5, t=4.567 df=4, *p=0.01, one-sample t test vs. 0%. ‘LFS’: 57.86 ± 12.4%, n=6, t=4.681 df=5, *p=0.005, one-sample t test vs. 0%. ‘LFS + LY367385, MPEP’: 22.18 ± 8.3%, n=6, t=2.663 df=5, *p=0.04, one-sample t test vs. 0%. One-way ANOVA: F=5.257, *p=0.0198. ‘Control’ vs. ‘LFS’: *p=0.0276; ‘LFS’ vs. ‘LFS + LY367385, MPEP’: *p=0.0309; Bonferroni’s multiple comparisons test.

**Figure 3.**
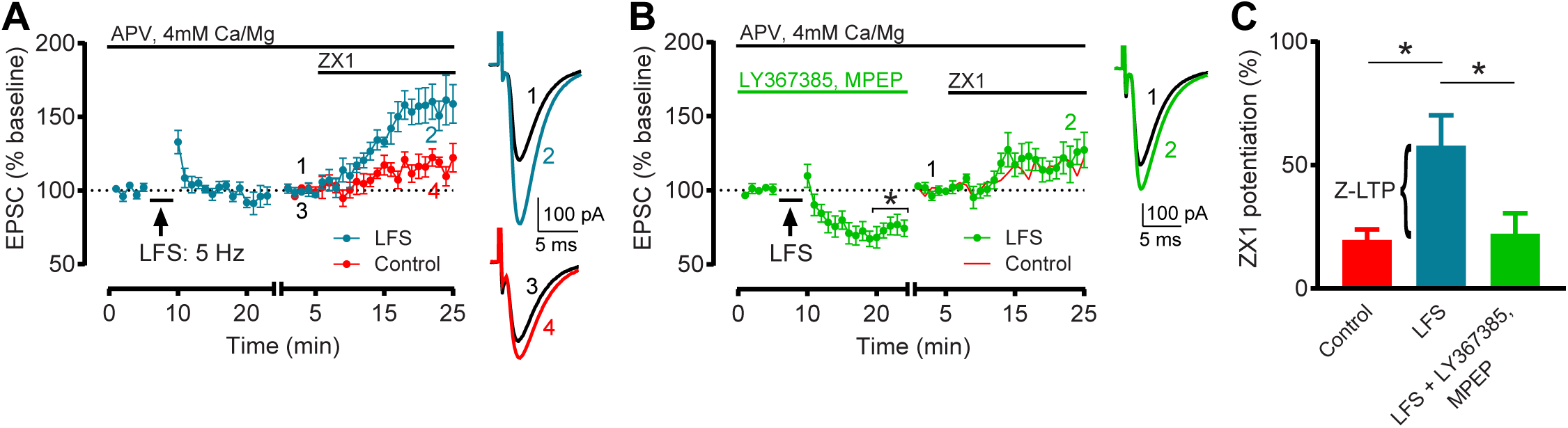
Low frequency stimulation (LFS) induces Z-LTP, which requires Group 1 mGluR activation. **(A)** Time course of AMPAR EPSC amplitude before and after LFS (5 Hz, 3 min), and before and after subsequent ZX1 application (cyan); and similar time course in interleaved control experiments (without LFS, red). **(B)** Same time course as in **A** but in the presence of LY367385 (100 μM) and MPEP (4 μM) (green). EPSC % baseline after LFS (mins. 20-24): n=6, *p=0.006, one-sample t test vs. 100%. Star (*) indicates significant LTD. Red line shows similar time course in controls replotted from **A**. For **(A-B)**, to examine the ZX1 potentiation after LFS, similar approach and renormalization as in **1C** was performed. Example traces show AMPAR EPSCs before and after ZX1. **(C)** Average ZX1 potentiation (% increase from baseline) during the last 5 min of ZX1 application. ‘Control’: n=5; ‘LFS’: n=6; ‘LFS + LY367385, MPEP’: n=6. LFS increased ZX1 potentiation compared to control; this increase was blocked by LY367385 and MPEP. One-way ANOVA/Bonferroni, *p=0.02. The increase in ZX1 potentiation is termed Z-LTP. Values represent mean ± SEM. Star (*) indicates p<0.05.

Figure 4: **(4A)** EPSC % baseline after 50 μM DHPG (average of mins. 16-20): 49.66 ± 5.1%, n=6, t=9.945 df=5, *p=0.0002, one-sample t test vs. 100%. One cell was included for analysis of EPSCs following 50 μM DHPG, but did not remain stable throughout subsequent ZX1 application and was excluded from analysis following ZX1 application, quantified in C. **(4B)** EPSC % baseline after 5 μM DHPG (average of mins. 16-20): 82.41 ± 7.4%, n=5, t=2.376 df=4, n.s. p=0.076, one-sample t test vs. 100%. **(4C)** ZX1 potentiation (%): ‘DHPG (50 μM)’: 93.51 ± 10.92%, n=5, t=8.561 df=4, *p=0.001, one-sample t test vs. 0%. ‘DHPG (5 μM)’: 0.44 ± 7.08%, n=5, t=0.06273 df=4, n.s. p=0.95, one-sample t test vs. 0%. One-way ANOVA: F=30.22, *p<0.0001. ‘Control’ vs. ‘DHPG (50 μM)’: *p<0.0001; ‘Control’ vs. ‘DHPG (5 μM)’: *p=0.01; Bonferroni’s multiple comparisons test. **(4D)** EPSC % baseline after 50 μM DHPG (average of mins. 16-20): 83.05 ± 5.9%, n=5, t=2.893 df=4, *p=0.044, one-sample t test vs. 100%. **(4E)** EPSC % baseline after LFS and 50 μM DHPG (average of mins. 20-24): 81.43 ± 9.1%, n=5, t=2.051 df=4, n.s. p=0.11, one-sample t test vs. 100%. **(4F)** ZX1 potentiation (%): ‘DHPG (50 μM)’: 55.83 ± 17.9%, n=5, t=3.12 df=4, *p=0.036, one-sample t test vs. 0%. ‘LFS + DHPG (50 μM)’: 74.65 ± 17.6%, n=5, t=4.246 df=4, *p=0.013, one-sample t test vs. 0%. One-way ANOVA: F=0.4126, n.s. p=0.6703. ‘LFS + DHPG (50 μM)’ vs. ‘LFS’: n.s. p=0.9181; ‘LFS + DHPG (50 μM)’ vs. ‘DHPG (50 μM)’: n.s. p=0.8553; Bonferroni’s multiple comparisons test.

**Figure 4.**
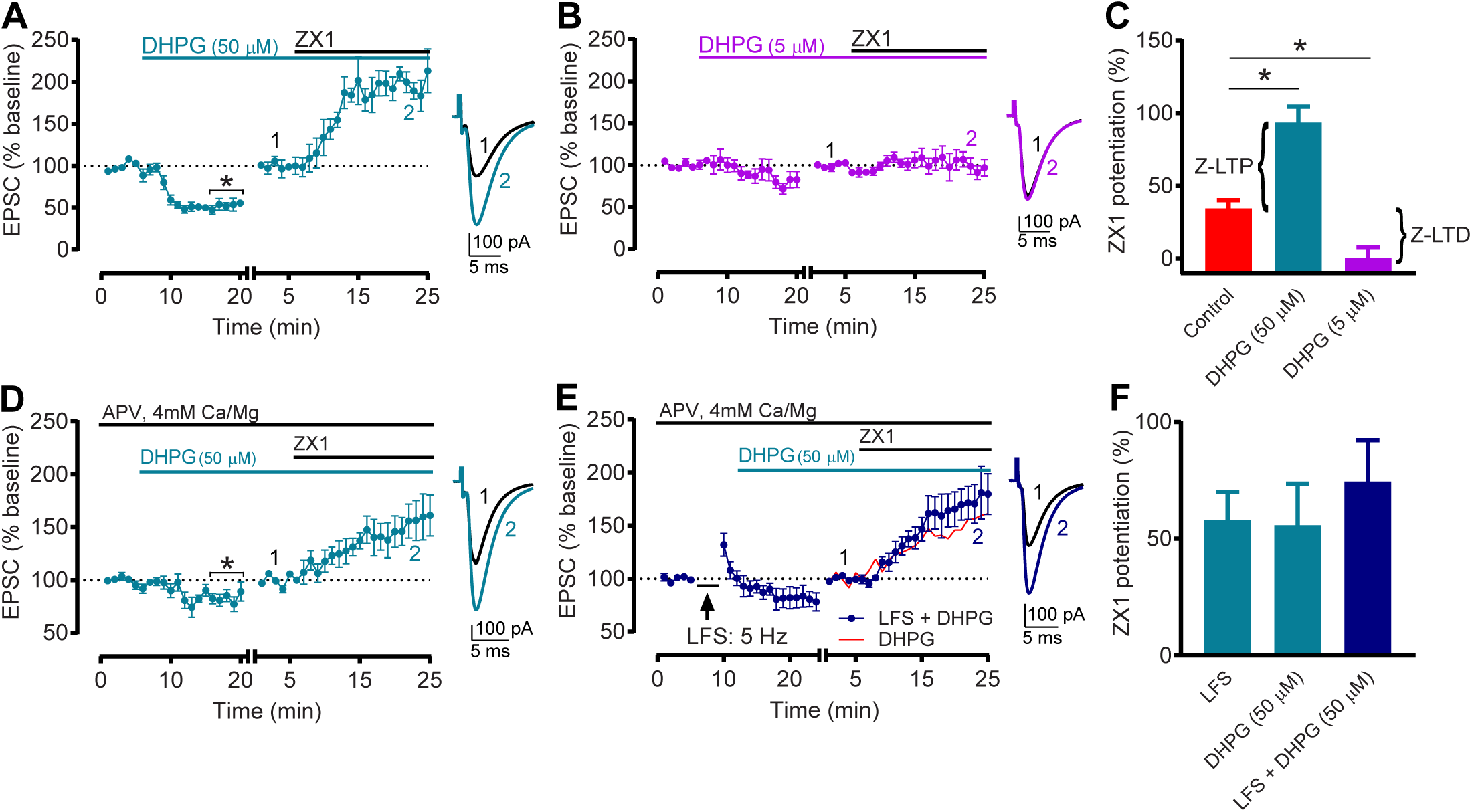
Group 1 mGluR activation is sufficient to induce Z-LTP and Z-LTD. **(A)** Time course of AMPAR EPSC amplitude before and after application of 50 μM DHPG, and before and after subsequent ZX1 application. EPSC % baseline after 50 μM DHPG (mins. 16-20): n=6, *p=0.0002, one-sample t test vs. 100%. Star (*) indicates significant synaptic depression. **(B)** Time course of AMPAR EPSC amplitude before and after application of 5 μM DHPG, and before and after subsequent ZX1 application. For **(A-B)**, to examine the ZX1 potentiation after DHPG application, after obtaining a stable baseline after DHPG, AMPAR EPSC amplitude was renormalized to the new baseline before ZX1 application. The renormalization is indicated by a gap and restart of timing in the x-axis. Example traces show AMPAR EPSCs before and after ZX1. **(C)** Average ZX1 potentiation (% increase from baseline) during the last 5 min of ZX1 application, with control data from **1D**. ‘DHPG (50 μM)’: n=5; ‘DHPG (5 μM)’: n=5. DHPG (50 μM) increased ZX1 potentiation compared to control, whereas DHPG (5 μM) reduced ZX1 potentiation compared to control. One-way ANOVA/Bonferroni, *p<0.0001. Increased and decreased ZX1 potentiation correspond to Z-LTP and Z-LTD, respectively. **(D)** Similar time course as in **A**, but in same extracellular conditions as in **3A-B**. EPSC % baseline after 50 μM DHPG (mins. 16-20): n=5, *p=0.044, one-sample t test vs. 100%. Star (*) indicates significant synaptic depression. **(E)** Time course of AMPAR EPSC amplitude before and after sequential LFS (5 Hz, 3 min) and application of 50 μM DHPG, and before and after subsequent ZX1 application, in same conditions as in **D**. For **(D-E)**, to examine the ZX1 potentiation, similar approach and renormalization as in **A-B** was performed. Example traces show AMPAR EPSCs before and after ZX1. **(F)** Average ZX1 potentiation (% increase from baseline) during the last 5 min of ZX1 application for the experiments in **D-E**, with LFS data from **3C**. ‘DHPG (50 μM)’: n=5; ‘LFS + DHPG (50 μM)’: n=5. Sequential LFS and DHPG (50 μM) did not increase ZX1 potentiation compared to LFS or DHPG (50 μM) alone. One-way ANOVA/Bonferroni: n.s. p=0.67. Values represent mean ± SEM. Star (*) indicates p<0.05.

Figure 5: **(5C)** DA-ZP1 fluorescence (% control): ‘+ DHPG (50 μM)’: 132.3 ± 9.096%, n=9,= *p=0.0039, Wilcoxon signed rank test vs. 100%. ‘+ DHPG (5 μM)’: 68.73 ± 11.99%, n=8, *p=0.0078, Wilcoxon signed rank test vs. 100%.

**Figure 5.**
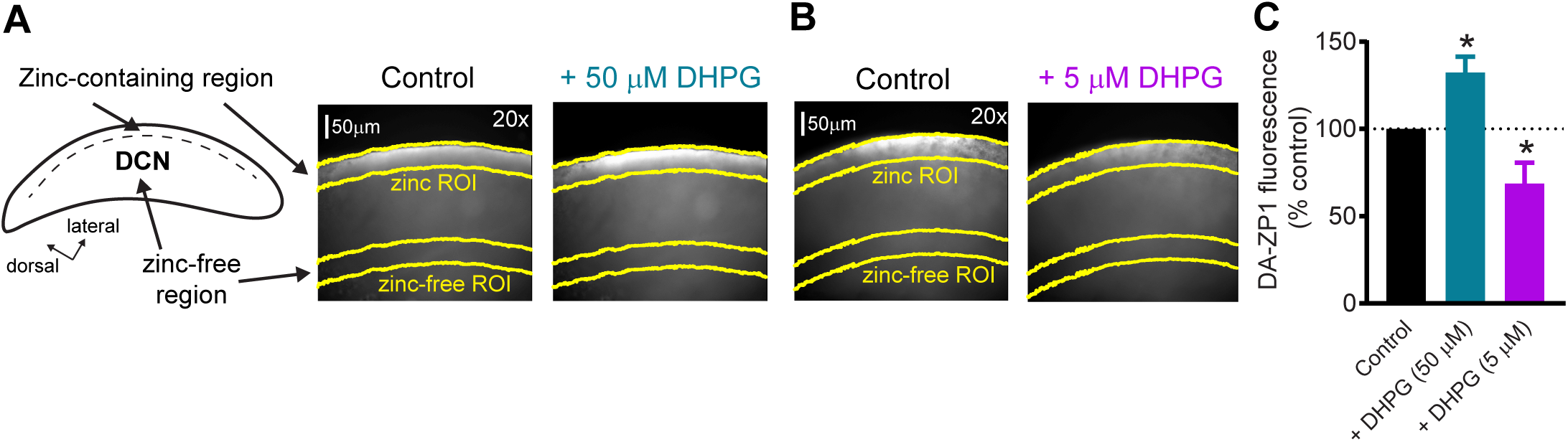
Group 1 mGluR activation bidirectionally modulates presynaptic zinc levels. **(A)** *Left:* Schematic of the DCN, showing the presynaptic zinc-containing region, and the zinc-free region. *Right:* 20x image of DA-ZP1 fluorescence, demonstrating the zinc-containing ROI (zinc ROI) and the zinc-free ROI, before and after application of 50 μM DHPG. **(B)** Same approach as in **A**, before and after application of 5 μM DHPG. **(C)** Average DA-ZP1 fluorescence after application of 50 μM or 5 μM DHPG, normalized to baseline fluorescence before DHPG application. ‘+ DHPG (50 μM)’: n=9, *p=0.004, Wilcoxon signed rank test vs. 100%. ‘+ DHPG (5 μM)’: n=8, *p=0.008, Wilcoxon signed rank test vs. 100%. Values represent mean ± SEM. Star (*) indicates p<0.05.

Figure 6: **(6C)** ZX1 potentiation (%): ‘Control’: 37.1 ± 3.1%, n=5, t=12.06 df=4, *p=0.0003, one-sample t test vs. 0%. ‘HFS’: 9.2 ± 5.2%, n=6, n.s. p=0.16, Wilcoxon signed rank test vs. 0%. ‘HFS + LY267385, MPEP’: 42.6 ± 8.2%, n=5, t=5.197 df=4, *p=0.007, one-sample t test vs. 0%. Kruskal-Wallis test: *p=0.0102. ‘Control’ vs. ‘HFS’: *p=0.0242; ‘HFS’ vs. ‘HFS + LY367385, MPEP’: *p=0.0428; Dunn’s multiple comparisons test. **(6D)** EPSC (% baseline): ‘Control’: 300nM: n=3, 90.68 ± 1.051%; 1μM: n=5, 70.89 ± 3.943%; 3μM: n=5, 54.61 ± 2.791%; 10μM: n=3, 40.72 ± 4.845%. ‘HFS’: 300nM: n=3, 90.24 ± 4.327%; 1μM: n=4, 69.52 ± 2.208%; 3μM: n=4, 52.1 ± 3.214%; 10μM: n=3, 37.89 ± 1.533%. Nonlinear fits: ‘Control’: Hill Slope=1.095, R^2^=0.9472. ‘HFS’: Hill Slope=1.128, R^2^=0.9765. **(6E)** IC_50_(μM): ‘Control’: 1.284 ± 0.3566. ‘HFS’: 1.267 ± 0.2321. Extra sum-of-squares F test: n.s. p=0.9687.

**Figure 6.**
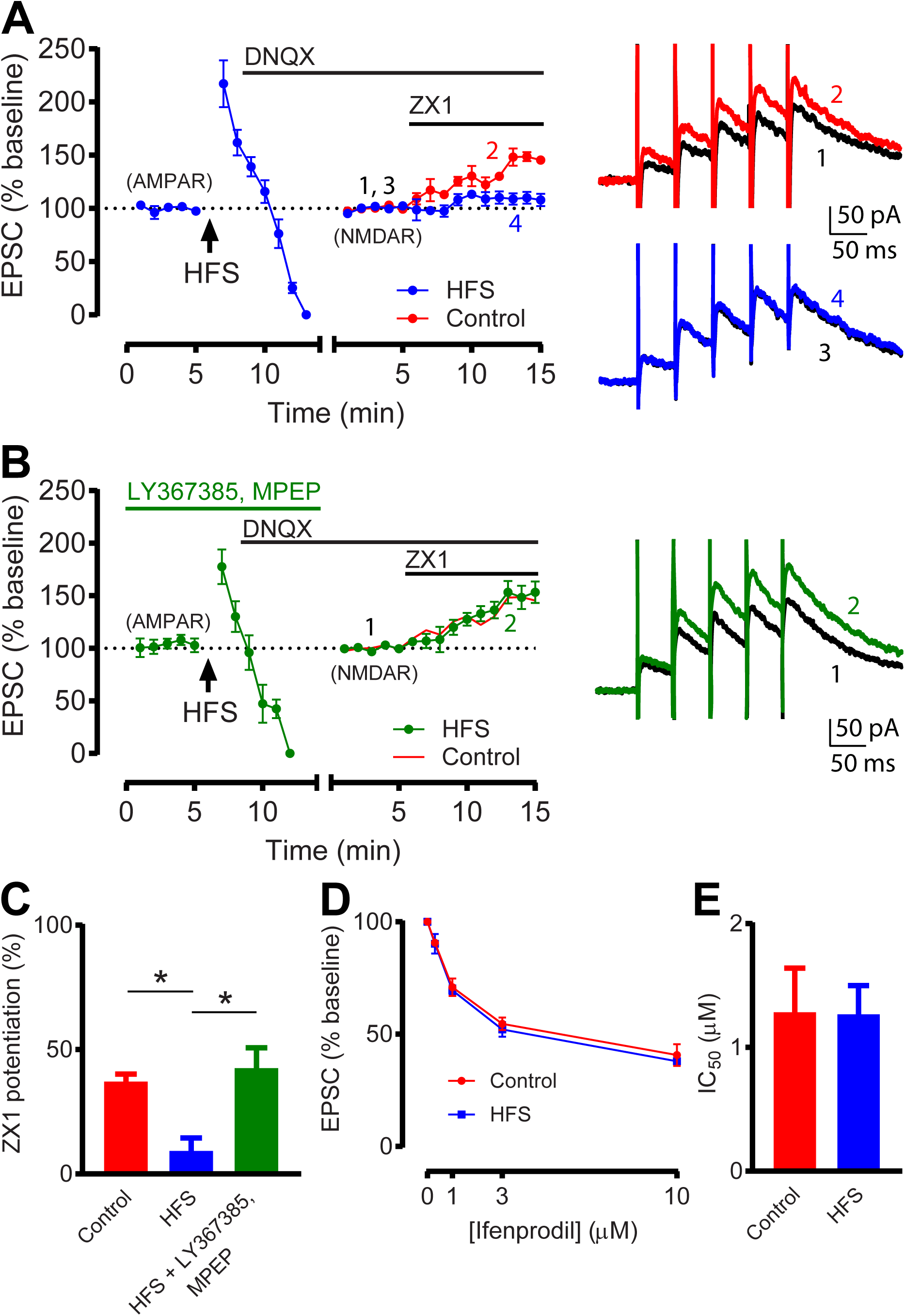
Group 1 mGluR-dependent Z-LTD reduces zinc inhibition of NMDARs. **(A)** *Left:* Time course of AMPAR EPSC amplitude before and after HFS, and NMDAR EPSC amplitude before and after subsequent ZX1 application (blue); and similar time course in interleaved control experiments (without HFS, red). *Right:* Example NMDAR EPSCs before and after ZX1 application. **(B)** *Left:* Same time course as in **A** but in the presence of LY367385 (100 μM) and MPEP (4 μM) (green). Red line shows controls replotted from **A**. *Right:* Example NMDAR EPSCs before and after ZX1 application. For **(A-B)**, after obtaining a stable baseline of AMPAR EPSCs, HFS was delivered, then DNQX (20 μM) was applied. NMDAR EPSCs were then recorded at +40 mV normalized to the baseline NMDAR EPSC amplitude before ZX1 application. The switch from AMPAR to NMDAR EPSC time course, and the renormalization of EPSC amplitude are indicated by a gap and restart of timing in the x-axis. **(C)** Average ZX1 potentiation (% increase from baseline) during the last 5 min of ZX1 application. ‘Control’: n=5; ‘HFS’: n=6; ‘HFS + LY367385, MPEP’: n=5. HFS reduced ZX1 potentiation compared to control; this reduction was blocked by LY367385 and MPEP. Kruskal-Wallis test/Dunn: *p=0.01. **(D)** Dose-response of NMDAR EPSCs (% baseline) for increasing concentrations of ifenprodil, in controls (red) and after HFS (blue). ‘Control’: n=3-5 per concentration; ‘HFS’: n=3-4 per concentration. IC_50_ of ifenprodil, from dose-responses in **D**. n.s. p=0.97, comparison of fits, extra sum-of-squares F test. Values represent mean ± SEM. Star (*) indicates p<0.05.

Figure 7: **(7B)** ZX1 potentiation (%): ‘N.E.’: 11.7 ± 8.56%, n=5, t=1.373 df=4, n.s. p=0.24, one-sample t test vs. 0%. ‘N.E. + AIDA’: 43.8 ± 8.05%, n=6, t=5.447 df=5, *p=0.003, one-sample t test vs. 0%. ‘N.E.’ vs. ‘N.E. + AIDA’: t=2.724 df=9, *p=0.024, unpaired t test. **(7C)** PPR: ‘N.E.’: 1.896 ± 0.19, n=5. ‘N.E. + AIDA: 2.056 ± 0.12, n=6. ‘N.E.’ vs. ‘N.E. + AIDA’: t=0.7446 df=9, n.s. p=0.476, unpaired t test. Normalized 1/CV^2^: ‘N.E. + AIDA’: 0.78 ± 0.22, n=6; n.s. p=0.44, Wilcoxon signed rank test vs. 1. **(7E)** ABR threshold (dB SPL): ‘Sham ipsi.’: 43.75 ± 3.24, n=8. ‘N.E. contra.’: 68.33 ± 3.07, n=6. ‘N.E. + AIDA contra.’: 65.71 ± 2.97, n=7. Kruskal-Wallis test: *p=0.0002. ‘Sham ipsi.’ vs. ‘N.E. contra.’: *p=0.0042; ‘Sham ipsi.’ vs. ‘N.E. + AIDA contra.’: *p=0.0076; ‘N.E. contra.’ vs. ‘N.E. + AIDA contra.’: n.s. p>0.9999; Dunn’s multiple comparisons test. ABR Wave I (μV): ‘Sham ipsi.’: 2.67 ± 0.31, n=8. ‘N.E. contra.’: 1.23 ± 0.13, n=6. ‘N.E. + AIDA contra.’: 1.25 ± 0.27, n=7. Kruskal-Wallis test: *p=0.0024. ‘Sham ipsi.’ vs. ‘N.E. contra.’: *p= 0.0387; ‘Sham ipsi.’ vs. ‘N.E. + AIDA contra.’: *p=0.0107; ‘N.E. contra.’ vs. ‘N.E. + AIDA contra.’: n.s. p>0.9999; Dunn’s multiple comparisons test.

**Figure 7.**
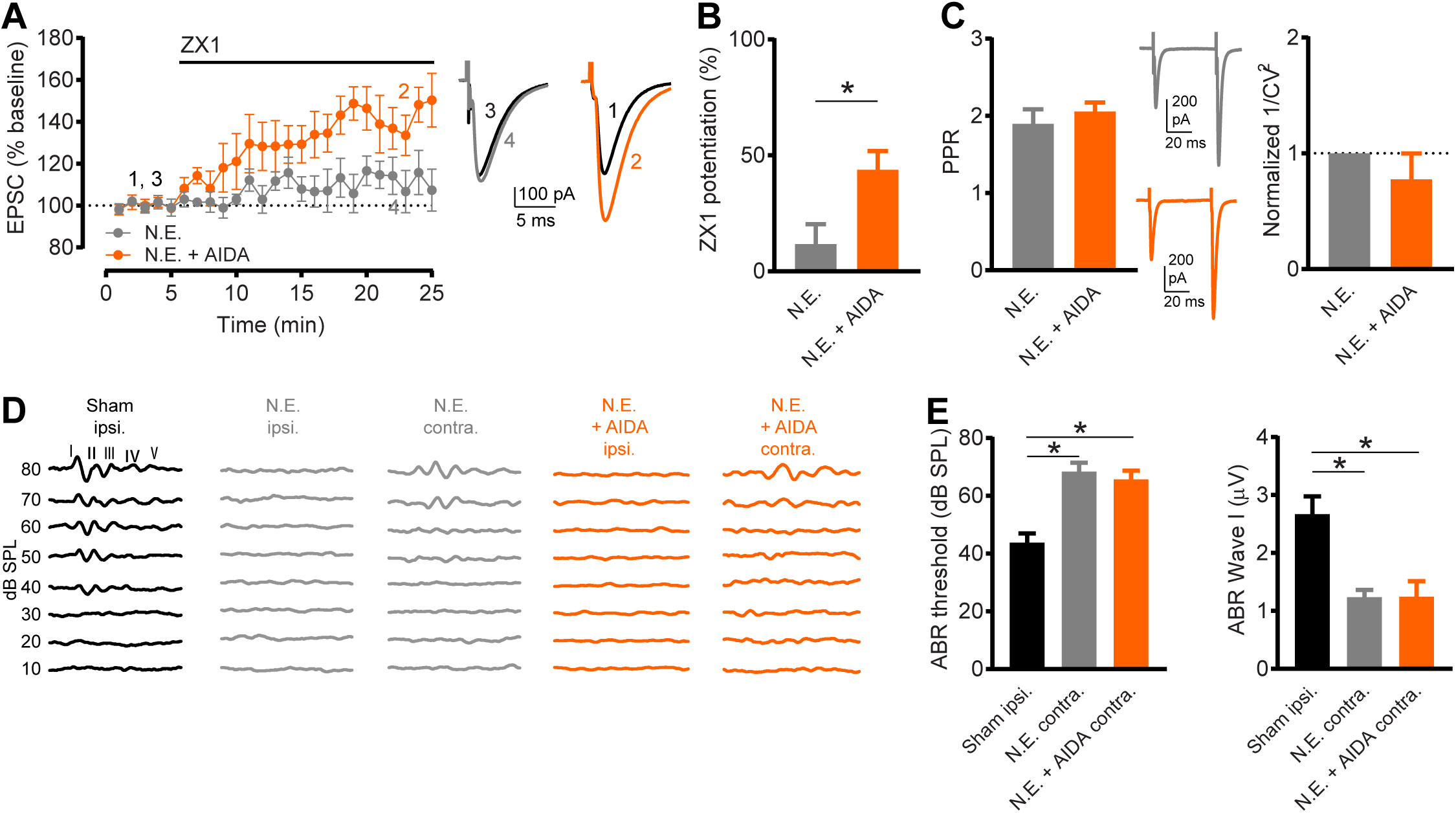
Sound-induced Z-LTD requires Group 1 mGluR activation. **(A)** Time course of AMPAR EPSC amplitude before and after ZX1 application in slices from N.E. mice (gray) and N.E. AIDA-treated mice (orange). Example traces show AMPAR EPSCs before and after ZX1. **(B)** Average ZX1 potentiation (% increase from baseline) during the last 5 min of ZX1 application. ‘N.E.’ (n=5) vs. ‘N.E. + AIDA’ (n=6): *p=0.024, unpaired t test. **(C)** *Left:* Average paired-pulse ratio (PPR, pulse 2 / pulse 1) of baseline AMPAR EPSCs in slices from N.E. mice and N.E. AIDA-treated mice. ‘N.E.’ (n=5) vs. ‘N.E. + AIDA’ (n=6): n.s. p=0.476, unpaired t test. Example traces show AMPAR EPSCs in response to two pulses. *Right:* coefficient of variation (CV) analysis (1/CV^2^) of baseline AMPAR EPSCs (pulse 1) in slices from N.E. mice and N.E. AIDA-treated mice, normalized to N.E. mice. ‘N.E. + AIDA’: n=6; n.s. p=0.44, Wilcoxon signed rank test vs. 1. **(D)** Example Auditory Brainstem Responses (ABRs, 10-80 dB SPL sound stimuli) from sham-exposed mice (recorded from sham-exposed, ipsilateral ear, black), N.E. mice (gray), and N.E. AIDA-treated mice (orange). Because no ABRs were detected in the ipsilateral ears of N.E. mice, ABRs were measured from ears contralateral to noise exposure. **(E)** *Left:* Average ABR thresholds (dB SPL). ‘Sham ipsi.’: n=8; ‘N.E. contra.’: n=6; ‘N.E. + AIDA contra.’: n=7. N.E. increased ABR thresholds compared to sham-exposed (*p=0.0002), but AIDA and N.E. did not affect increases in ABR thresholds compared to N.E. alone, Kruskal-Wallis test/Dunn. *Right:* Average ABR Wave I amplitude (μV). ‘Sham ipsi.’: n=8; ‘N.E. contra.’: n=6; ‘N.E. + AIDA contra.’: n=7. N.E. decreased ABR Wave I amplitude compared to sham-exposed (*p=0.0024), but AIDA and N.E. did not affect decreases in ABR Wave I amplitude compared to N.E. alone, Kruskal-Wallis test/Dunn. Values represent mean ± SEM. Star (*) indicates p<0.05.

## Results

### Bidirectional activity-dependent long-term synaptic zinc plasticity requires Group 1 mGluR activation

To investigate the mechanisms underlying synaptic zinc plasticity, we first determined whether we could induce long-term synaptic zinc plasticity in DCN PF synapses in mouse brain slices. In these synapses, synaptic zinc inhibits AMPAR and NMDAR EPSCs via postsynaptic mechanisms. This has been evidenced by application of ZX1, a fast, high-affinity extracellular zinc chelator, which potentiates AMPAR and NMDAR EPSCs (Anderson *et al.*, 2015; Kalappa *et al.*, 2015; Kalappa & Tzounopoulos, 2017). This *ZX1 potentiation* of AMPA and NMDAR EPSCs is dependent on ZnT3, the transporter that loads zinc into synaptic vesicles (Palmiter *et al.*, 1996; Cole *et al.*, 1999). Moreover, reductions in ZX1 potentiation reflect reductions in synaptic zinc levels and release (Kalappa *et al.*, 2015). Therefore, in this study, we used the amount of ZX1 potentiation of AMPAR and NMDAR EPSCs to monitor synaptic zinc signaling, and long-term synaptic zinc plasticity, in DCN PF synapses.

Consistent with previous studies, we found that ZX1 potentiated postsynaptic PF AMPAR EPSCs in DCN cartwheel cells (CWCs), a class of inhibitory interneurons (Figure 1 A-B) (Kalappa *et al.*, 2015; Kalappa & Tzounopoulos, 2017). We then tested whether we can induce long-term synaptic zinc plasticity by using patterns of synaptic activation that induce long-term plasticity of glutamatergic synaptic strength in DCN PF synapses, such as LTP and LTD (Fujino & Oertel, 2003; Tzounopoulos *et al.*, 2004; Tzounopoulos *et al.*, 2007). We started by examining the effect of ZX1 on AMPAR EPSCs following high-frequency stimulation of PFs (HFS, 3 × 100 Hz for 1 sec, 10 sec inter-stimulus interval), which induces LTP (Fujino & Oertel, 2003). After applying HFS and inducing LTP (Figure 1 C), we renormalized AMPAR EPSC amplitude to quantify the amount of ZX1 potentiation. (Figure 1 C). After HFS, ZX1 application did not potentiate AMPAR EPSCs (Figure 1 C-D). The loss of ZX1 potentiation indicates a loss of zinc-mediated inhibition of AMPARs, suggesting that HFS caused a long-term reduction in synaptic zinc signaling, termed *Z-LTD* (Figure 1 D). Furthermore, these results suggest that Z-LTD, by reducing zinc-mediated inhibition of AMPAR EPSCs and thus enhancing baseline synaptic strength, is a new mechanism of HFS-induced LTP. For a discussion on the impact of synaptic zinc plasticity in the context of other long-term plasticity mechanisms, see *Discussion* section, *Implications of Z-LTP and Z-LTD for LTD and LTP*.

After establishing that HFS caused Z-LTD, we then studied the underlying mechanisms. NMDARs contribute to the induction of LTP and LTD in the DCN and most central synapses (Malenka & Nicoll, 1993; Fujino & Oertel, 2003; Tzounopoulos *et al.*, 2004; Tzounopoulos *et al.*, 2007). To test the role of NMDARs in the induction of Z-LTD, we blocked NMDARs with APV (NMDAR antagonist, 50 μM; Figure 2 A). As evidenced by the lack of ZX1 potentiation of AMPAR EPSCs after HFS, APV did not affect Z-LTD (Figure 2 A, C), indicating that NMDARs are not required for the induction of Z-LTD.

Parallel fiber synapses in the DCN also exhibit glutamatergic plasticity that involves metabotropic glutamate receptor (mGluR) signaling (Fujino & Oertel, 2003). Furthermore, Group 1 (G1) mGluRs are expressed in CWCs and in the DCN molecular layer, where PF terminals reside (Wright *et al.*, 1996; Bilak & Morest, 1998). We therefore tested whether G1 mGluR activation is necessary for Z-LTD. To test this hypothesis, we repeated the experiment shown in Figure 2A, but we now blocked G1 mGluRs with MPEP (4 μM, mGluR5-selective antagonist) and LY367385 (100 μM, mGluR1-selective antagonist) (Figure 2 B). Under these conditions, ZX1 potentiation was observed after HFS, indicating that HFS did not induce Z-LTD (Figure 2 B-C). This result demonstrates that G1 mGluR activation is necessary for the induction of Z-LTD.

Glutamatergic plasticity is bidirectional: synapses undergo LTP or LTD in response to high-or low-frequency stimulation, respectively (Mulkey & Malenka, 1992; Malenka & Nicoll, 1993; Fujino & Oertel, 2003). To determine whether long-term synaptic zinc plasticity is bidirectional, we tested whether low-frequency stimulation (LFS) increases zinc signaling (Figure 3 A). Because the induction of zinc plasticity depends on mGluR activation, we used conditions that favor mGluR-dependent LTD, such as LFS (5 Hz, 3 min), blockade of NMDARs with APV, and high extracellular concentrations of divalent ions (4 mM Ca^2+^ and Mg^2+^) (Oliet *et al.*, 1997). Compared to interleaved control experiments, LFS increased the amount of subsequent ZX1 potentiation (Figure 3 A, C). Increased ZX1 potentiation indicates increased zinc-mediated inhibition of AMPARs, suggesting that LFS caused a long-term increase in synaptic zinc signaling, termed *Z-LTP* (Figure 3 C). By enhancing zinc-mediated inhibition of AMPAR EPSCs, Z-LTP is a new mechanism of LFS-induced LTD.

Note that control ZX1 potentiation in these conditions (Figure 3 A, C) was slightly less, albeit not significantly different (*p*=0.11, unpaired *t* test), than previous control experiments performed in ACSF with 2.4/1.3 mM of extracellular Ca^2+^/Mg^2+^ (Figure 1 D). This is likely due to reduced neuronal excitability in higher divalent concentrations (Oliet *et al.*, 1997; Kalappa *et al.*, 2015). Together, these results show that LFS induced Z-LTP, thus demonstrating that activity-dependent plasticity of zinc signaling is bidirectional: HFS induces long-term depression of zinc signaling (Z-LTD), whereas LFS induces long-term potentiation of zinc signaling (Z-LTP).

We next tested whether G1 mGluR activation is necessary for the induction of Z-LTP. In the presence of MPEP and LY367385, LFS did not increase the amount of ZX1 potentiation (Figure 3 B-C), indicating that G1 mGluR activation is necessary for the induction of Z-LTP. Together, these results reveal that activation of G1 mGluR signaling is necessary for the induction of Z-LTP and Z-LTD.

### Group 1 mGluR activation is sufficient to induce bidirectional long-term synaptic zinc plasticity

Is activation of G1 mGluRs sufficient to induce Z-LTP and Z-LTD? Because G1 mGluRs are required for both increases and decreases in synaptic zinc signaling by different stimulation paradigms, we hypothesized that the direction of plasticity depends on the differential activation of G1 mGluRs during HFS and LFS. To test this, we applied high or low concentrations of DHPG (G1 mGluR agonist, 50 μM or 5 μM). Consistent with previous studies, application of 50 μM DHPG caused a significant depression of synaptic strength (Figure 4 A) (Huber *et al.*, 2001; Snyder *et al.*, 2001; Wisniewski & Car, 2002). After applying 50 μM DHPG, obtaining a new stable baseline, and then applying ZX1, we observed that the ZX1 potentiation of EPSCs was significantly increased compared to control experiments (Figure 4 A, C). This result indicates that a high concentration of DHPG increases synaptic zinc signaling: G1 mGluR activation is sufficient to induce Z-LTP.

Because Z-LTP and Z-LTD induced by LFS and HFS depend on G1 mGluR activation (Figure 2 and 3), we next tested whether application of a lower concentration of DHPG causes Z-LTD. After applying 5 μM DHPG and obtaining a new stable baseline, ZX1 did not potentiate EPSCs, consistent with Z-LTD induction (Figure 4 B-C). Together, these results demonstrate that G1 mGluR activation is sufficient to cause bidirectional zinc plasticity. Furthermore, the direction of zinc plasticity depends on the concentration of DHPG: 50 μM DHPG causes Z-LTP, whereas 5 μM DHPG causes Z-LTD (Figure 4 C). These results are consistent with the notion that bidirectional zinc plasticity depends on differential activation of G1 mGluRs by either LFS/HFS or high/low concentrations of DHPG.

Electrical synaptic stimulation with LFS/HFS or pharmacological activation of G1 mGluRs with high/low concentrations of DHPG induce bidirectional synaptic zinc plasticity; however, it is unknown whether these two different methods induce mechanistically similar synaptic zinc plasticity. To explore this, we compared the amount of Z-LTP elicited by applying sequential LFS and 50 μM DHPG to the amount of Z-LTP elicited by LFS or 50 μM DHPG alone. If electrical and pharmacological manipulations induce Z-LTP by different mechanisms, then LFS and 50 μM DHPG application should yield an additive effect on Z-LTP, and subsequent ZX1 potentiation should be greater than that following LFS alone or application of 50 μM DHPG alone. To test this, we performed interleaved experiments to determine the effect of 50 μM DHPG alone, under the conditions used for LFS-induced Z-LTP as in Figure 3, with experiments involving stimulation with LFS and subsequent DHPG application (Figure 4 D-E). Under these conditions, ZX1 potentiation following application of 50 μM DHPG was similar to ZX1 potentiation following LFS (Figure 4 F). Importantly, ZX1 potentiation after sequential LFS and 50 μM DHPG was not significantly greater than ZX1 potentiation after LFS or DHPG alone (Figure 4 F). Together, these results show that LFS occluded the effect of 50 μM DHPG; thus, LFS and DHPG induce Z-LTP likely via a common mechanistic pathway.

### Group 1 mGluR activation bidirectionally modulates presynaptic zinc levels

We used activity-dependent changes in the amount of ZX1 potentiation of AMPAR EPSCs for assessing changes in synaptic zinc signaling (Z-LTP and Z-LTD). However, ZX1 potentiation is determined by the postsynaptic zinc-mediated inhibition of AMPAR EPSCs, as well as the amount of presynaptic zinc release (Kalappa *et al.*, 2015). Because previous studies demonstrated sensory experience-dependent, long-term modulation of presynaptic zinc levels (Nakashima & Dyck, 2009; Kalappa *et al.*, 2015), we hypothesized that Z-LTP and Z-LTD are expressed, at least in part, by the modulation of presynaptic zinc levels. To quantify potential changes in presynaptic zinc levels, we used DA-ZP1, a fluorescent intracellular zinc sensor capable of tracking presynaptic zinc levels in PF terminals (Kalappa *et al.*, 2015; Zastrow *et al.*, 2016). DA-ZP1 produces a band of fluorescence within the DCN molecular layer in wild type mice. This fluorescent signal is absent in mice lacking the vesicular ZnT3 transporter, thus demonstrating that the signal is due to ZnT3-dependent, synaptic zinc (Kalappa *et al.*, 2015; Zastrow *et al.*, 2016). To induce Z-LTP and Z-LTD, we applied DHPG, which is mechanistically similar to electrically-induced Z-LTP and Z-LTD (Figure 4 F) and capable of inducing robust synaptic zinc plasticity in many terminals in the slice. To test for changes in presynaptic zinc levels, we imaged DA-ZP1 fluorescence in the same region of the same DCN slice before and after DHPG application (50 μM or 5 μM) (Figure 5 A; see *Materials and Methods*). Application of 50 μM DHPG increased DA-ZP1 fluorescence, indicating increased presynaptic zinc levels in PF terminals, which is consistent with Z-LTP (Figure 5 A, C). In contrast, application of 5 μM DHPG reduced DA-ZP1 fluorescence, indicating reduced zinc levels, which is consistent with Z-LTD (Figure 5 B-C).

Together, these results demonstrate that differential activation of G1 mGluRs, by application of different concentrations of DHPG, causes bidirectional modulation of presynaptic zinc levels. Furthermore, these results are consistent with our electrophysiological experiments: 50 μM DHPG results in Z-LTP by increasing presynaptic zinc levels, whereas 5 μM DHPG results in Z-LTD by reducing presynaptic zinc levels. Although these results do not rule out potential postsynaptic mechanisms of Z-LTP and Z-LTD, they demonstrate that Z-LTP and Z-LTD are associated with modulation of presynaptic zinc levels.

### G1 mGluR-dependent Z-LTD reduces zinc-mediated inhibition of NMDARs

Z-LTP and Z-LTD involve modulation of presynaptic zinc signaling (Figure 5). Based on this finding, the induction of long-term synaptic zinc plasticity should also affect postsynaptic NMDAR EPSCs, which are inhibited by zinc via direct high-affinity NMDAR allosteric modulation (Paoletti *et al.*, 1997; Vergnano *et al.*, 2014). To test this prediction, we quantified the ZX1 potentiation of NMDAR EPSCs after inducing Z-LTD with HFS. To monitor NMDAR EPSCs, we used a short train of presynaptic stimulation (5 pulses at 20 Hz) to activate extrasynaptic NMDARs, for NMDAR EPSCs recorded in somata of CWCs are mostly mediated by extrasynaptic NMDARs activated by glutamate spillover during this short train (Anderson *et al.*, 2015). To avoid keeping CWCs at +40 mV for too long while recording NMDAR EPSCs, and to maintain the same induction protocol used in our previous experiments, we initially recorded AMPAR EPSCs at −70 mV and then applied HFS (Figure 6 A). Subsequently, we blocked AMPARs with DNQX (20 μM, AMPA/kainate receptor antagonist) and recorded at +40 mV to obtain a stable baseline of NMDAR EPSCs before applying ZX1 (Figure 6 A). Consistent with our results on AMPAR EPSCs, after HFS, ZX1 no longer potentiated NMDAR EPSCs, whereas ZX1 potentiated NMDAR EPSCs in interleaved control experiments, where HFS was not applied (Figure 6 A, C). These results demonstrate that Z-LTD reduces zinc-mediated inhibition of NMDARs. To determine whether this plasticity shares the same mechanism as Z-LTD evidenced by changes in the ZX1 potentiation of AMPAR EPSCs, we tested whether G1 mGluR signaling is required. Indeed, application of MPEP and LY367385 blocked the observed Z-LTD, evidenced by the ZX1 potentiation of NMDAR EPSCs after HFS (Figure 6 B-C). Together, our results suggest that G1 mGluR-dependent synaptic zinc plasticity modulates zinc-mediated inhibition of AMPARs and NMDARs similarly, suggesting that it is independent of the mode of action of synaptic zinc on its postsynaptic targets. This supports our findings that zinc plasticity is expressed, at least in part, by changes in presynaptic zinc levels.

However, the contribution of postsynaptic mechanisms in synaptic zinc plasticity cannot be excluded. To address this possibility, we tested whether activity-dependent changes in postsynaptic NMDAR subunit composition could modulate zinc sensitivity. NMDARs are composed of two GluN1 subunits and two GluN2 subunits (Traynelis *et al.*, 2010). GluN2A-containing NMDARs (GluN1/GluN2A diheteromers and GluN2/GluN2A/GluN2B triheteromers) have nanomolar affinity for zinc, whereas GluN1/GluN2B diheteromers have micromolar affinity (Paoletti *et al.*, 1997; Rachline *et al.*, 2005; Tovar & Westbrook, 2012; Hansen *et al.*, 2014). Therefore, the reduced zinc-mediated inhibition of NMDAR EPSCs after HFS, evidenced by reduced ZX1 potentiation (Figure 6 A), could be explained by an increase in the proportion of GluN2B subunits. We therefore tested whether HFS increases the sensitivity of NMDAR EPSCs to ifenprodil, a GluN2B-selective antagonist (Figure 6 D-E) (Tovar & Westbrook, 2012; Hansen *et al.*, 2014). Compared to controls, HFS did not affect the ifenprodil sensitivity (IC_50_) of NMDAR EPSCs (Figure 6 D-E). This indicates that HFS-induced plasticity does not alter the proportions of GluN2B vs. GluN2A NMDAR subunits, suggesting that Z-LTD is not due to reduced zinc sensitivity caused by a decrease in the relative contribution of GluN2A vs. GluN2B in the NMDAR EPSC. Therefore, these results further support that zinc plasticity is expressed by changes in presynaptic zinc levels, rather than postsynaptic receptor modifications.

### Sound-induced zinc plasticity requires Group 1 mGluRs *in vivo*

Our experiments described here, using *in vitro* brain slice electrophysiology in the DCN point toward a mechanism of bidirectional long-term synaptic zinc plasticity dependent on G1 mGluR activation. We therefore hypothesized that G1 mGluR activation may also be necessary for the reduction in synaptic zinc signaling observed in the DCN after sound exposure (Kalappa *et al.*, 2015). To test this hypothesis, we quantified the ZX1 potentiation of PF EPSCs in DCN slices from mice exposed to loud sound (116 dB, 4 hours). Consistent with sound-induced LTD and previous studies (Kalappa *et al.*, 2015), we did not observe ZX1 potentiation in slices from noise-exposed (N.E.) mice (Figure 7 A-B). To test whether G1 mGluRs are necessary for the reduced zinc signaling in slices from N.E. mice, we administered a systemic, blood brain barrier-permeable G1 mGluR antagonist (AIDA, i.p., 2 mg/kg; twice: 30 min before and 1.5 hours after beginning the noise exposure). Indeed, we observed ZX1 potentiation in slices from N.E. mice treated with AIDA (Figure 7 A-B), suggesting that *in vivo* inhibition of G1 mGluR activity blocked the sound-induced Z-LTD.

Although AIDA treatment blocked Z-LTD in DCN PF synapses (Figure 7 A-B), it did not affect assays that are sensitive to presynaptic glutamate release probability, such as paired-pulse ratio (PPR) and coefficient of variation (CV) analysis (Figure 7 C). This indicates that sound-induced G1 mGluR-dependent Z-LTD specifically modulates synaptic zinc signaling, without affecting presynaptic glutamate signaling in PFs. Furthermore, AIDA treatment did not affect sound-induced hearing loss in N.E. mice, quantified with Auditory Brainstem Responses (ABRs) (Figure 7 D). ABRs reflect the synchronous activity, arising from the auditory nerve (Wave I), of auditory brainstem nuclei to the inferior colliculus (Waves II-V) in response to sound stimuli. Elevated ABR thresholds indicate increased hearing thresholds. However, similar ABR thresholds may be accompanied by differences in the suprathreshold response of Wave I, which could reflect differential degeneration of the auditory nerve (Kujawa & Liberman, 2009). AIDA treatment did not affect noise-induced changes in either ABR thresholds or Wave I amplitude (Figure 7 E), thus indicating that the effect of AIDA on blocking Z-LTD is not due to differential noise-induced hearing loss after AIDA treatment. Together, these results demonstrate that sound-induced Z-LTD requires G1 mGluR activation, consistent with our *in vitro* results.

## Discussion

Our results show that long-term synaptic zinc plasticity is an experience-, G1 mGluR-dependent mechanism that bidirectionally modulates synaptic zinc signaling in the DCN. Is this a general mechanism that applies to all synaptic zinc-containing brain areas? Synaptic zinc is present throughout the neocortex and other brain structures, such as the amygdala and the hippocampus (McAllister & Dyck, 2017). Moreover, synaptic zinc is modulated by sensory activity throughout the sensory cortex (McAllister & Dyck, 2017), shapes the gain of central sensory responses (Anderson *et al.*, 2017), and when upregulated by optic nerve injury, it inhibits retinal ganglion cell survival and axon regeneration (Li *et al.*, 2017). It is therefore likely, although not tested here, that the reported long-term synaptic zinc plasticity mechanism is a general mechanism that dynamically modulates sensory processing for adaptation to different sensory environments and injury.

Whereas the exact synaptic, natural, and ethologically relevant stimuli that elicit Z-LTP and Z-LTD remain unknown, here we developed *in vitro* and *in vivo* models for studying Z-LTP and Z-LTD. This is a crucial step towards further elucidation of the detailed natural stimuli eliciting long-term synaptic zinc plasticity, as well as the precise cellular and molecular mechanisms underlying the induction and expression of Z-LTP and Z-LTD. Moreover, our model will be useful for probing the unknown behavioral consequences of Z-LTP and Z-LTD.

### Mechanisms of Group 1 mGluR-dependent Z-LTP and Z-LTD

Our results show that differential activation of G1 mGluRs, by either LFS/HFS or high/low concentrations of DHPG, determines the induction and direction of long-term synaptic zinc plasticity. Prolonged LFS causes Z-LTP, similarly to G1 mGluR activation with 50 μM DHPG; whereas, brief HFS causes Z-LTD, similarly to activation with 5 μM DHPG. This suggests that prolonged LFS activates G1 mGluR signaling differently than brief HFS.

Group 1 mGluRs, mGluR1 and mGluR5, are linked to the IP_3_-Diacylglycerol (DAG) signaling pathway, leading to intracellular rises in Ca^2+^ from intracellular stores (Abdul-Ghani *et al.*, 1996; Conn & Pin, 1997; Kim *et al.*, 2008). In the hippocampus, LFS induces G1 mGluR-mediated LTD via postsynaptic AMPAR endocytosis involving Ca^2+^ release from endoplasmic reticulum (ER) stores and dendritic protein synthesis (Huber *et al.*, 2000; Holbro *et al.*, 2009; Luscher & Huber, 2010; Pick & Ziff, 2018). Moreover, in the hippocampus, HFS or theta-burst stimulation induces G1 mGluR-mediated LTP, also involving ER Ca^2+^ release, resulting in postsynaptic AMPAR/NMDAR trafficking or enhanced presynaptic glutamate release (Topolnik *et al.*, 2006; Wu *et al.*, 2008; Anwyl, 2009). It remains unknown whether G1 mGluR-dependent Z-LTP and Z-LTD are downstream effects of the same signaling pathways that induce LTD and LTP, or occur through separate mechanisms. Nonetheless, we propose, albeit not tested here, that differential G1 mGluR activation, by LFS/HFS, leads to subsequent release of different amounts or types of intracellular Ca^2+^ signals. Different Ca^2+^ signals may in turn activate diverse signaling pathways that ultimately lead to increased and decreased synaptic zinc signaling. An analogue that comes to mind is the mechanism via which differential activation of NMDARs, by various levels of synaptic activity, leads to variable Ca^2+^ levels and signaling, ultimately determining the induction of both LTP and LTD (Malenka & Bear, 2004).

Our results suggest that increases or decreases in synaptic zinc signaling, evidenced by increased or decreased ZX1 potentiation of EPSCs, are mediated by bidirectional modulation of vesicular zinc levels and subsequent synaptic zinc release. High or low concentrations of DHPG, which induce Z-LTP and Z-LTD, increase or decrease presynaptic zinc levels in PF terminals (Figure 5). Furthermore, synaptic zinc plasticity modulates zinc-mediated inhibition of NMDARs as well as AMPARs, and this effect on NMDARs cannot be explained by postsynaptic changes in the relative contributions of GluN2A vs. GluN2B subunits in the NMDAR EPSCs (Figure 6). Although we cannot fully exclude potential contributions of postsynaptic mechanisms in synaptic zinc plasticity, our results support that synaptic zinc plasticity is mainly mediated by activity-dependent modulation of presynaptic zinc levels and signaling, and are consistent with previous studies demonstrating experience-dependent modulation of vesicular zinc levels in the somatosensory cortex (Brown & Dyck, 2002, 2005), visual cortex (Dyck *et al.*, 2003), optic nerve (Li *et al.*, 2017), and the DCN (Kalappa *et al.*, 2015).

Cartwheel cells express G1 mGluRs, particularly mGluR1, suggesting that the locus of induction of zinc plasticity is postsynaptic (Wright *et al.*, 1996). Because Z-LTP and Z-LTD involve modulation of presynaptic zinc levels, one suggestion is the presence of a retrograde signal from CWCs involved in the expression of Z-LTP and Z-LTD in PFs. Alternatively, the presence of mGluR1 on axon terminals in the DCN molecular layer may support a presynaptic locus of induction (Bilak & Morest, 1998). Because ZnT3 determines vesicular zinc levels (Palmiter *et al.*, 1996; Cole *et al.*, 1999), modulation of ZnT3 expression or function may underlie the expression of Z-LTP and Z-LTD. In the retina, optic nerve injury increases ZnT3 immunostaining, supporting that increases in ZnT3 expression mediate increases in synaptic zinc levels (Li *et al.*, 2017). However, in the barrel cortex, whisker plucking increases the vesicular zinc content and the density of zinc-containing synapses, but does not alter either ZnT3 protein or mRNA levels (Brown & Dyck, 2002; Liguz-Lecznar *et al.*, 2005; Nakashima & Dyck, 2010; Nakashima *et al.*, 2011). Furthermore, in barrel cortical layers IV and V, the density of excitatory synapses remains unchanged despite the increased density of zinc-containing synapses, indicating that some previously excitatory non-zinc-containing synapses were converted to zinc-containing synapses (Nakashima & Dyck, 2010). Together, these studies suggest that changes in vesicular zinc content can occur without affecting glutamatergic synapses, likely via functional modulation of pre-existing ZnT3. This may also explain our electrophysiological results after sound exposure, because sound exposure caused Z-LTD without affecting presynaptic glutamate dynamics (Figure 7 C). In the context of ZnT3 modulation, it is interesting that the vesicular glutamate transporter 1 (VGlut1), which is co-targeted to synaptic vesicles with ZnT3, increases ZnT3 zinc transport in cultured cells (Salazar *et al.*, 2005). Because VGlut1 is highly expressed in the DCN molecular layer (Zhou *et al.*, 2007), one hypothesis is that modulation of VGlut1 may modulate ZnT3 function in PF terminals. However, the independent modulation of presynaptic glutamate and zinc dynamics after sound exposure (Figure 7 C) suggests a VGlut1-independent mechanism of ZnT3 modulation. While our results reveal a role for G1 mGluRs in Z-LTP and Z-LTD, future experiments will be necessary to determine the detailed induction and expression mechanisms.

### Implications of Z-LTP and Z-LTD for short-term plasticity

Previous studies in DCN PF synapses revealed that synaptic zinc triggers endocannabinoid synthesis, which inhibits presynaptic glutamate release and modulates short-term plasticity (Perez-Rosello *et al.*, 2013; Kalappa & Tzounopoulos, 2017). During high-frequency (50 Hz) trains, synaptic zinc inhibits AMPAR EPSCs during the first few stimuli, but enhances steady-state EPSCs in subsequent stimuli by recruiting endocannabinoid signaling and enhancing synaptic facilitation (Kalappa & Tzounopoulos, 2017). Therefore, long-term increases in zinc signaling, via Z-LTP, would enhance endocannabinoid activation during subsequent stimulus trains, increase synaptic facilitation, and further enhance steady-state EPSCs. Conversely, long-term decreases in zinc signaling, via Z-LTD, would reduce endocannabinoid activation, decrease synaptic facilitation, and suppress steady-state EPSCs.

Following stimulus trains, zinc-mediated endocannabinoid activation causes short-term depression and inhibits short-term facilitation (Perez-Rosello *et al.*, 2013). Therefore, Z-LTP and Z-LTD are expected to shift the balance between short-term facilitation and short-term depression in DCN synapses. Z-LTP will enhance subsequent zinc-mediated short-term depression, whereas Z-LTD will enhance short-term facilitation. Taken together, our results highlight a powerful mechanism by which long-term bidirectional zinc plasticity may modulate short-term glutamatergic synaptic plasticity.

### Implications of Z-LTP and Z-LTD for LTD and LTP

In central synapses, including DCN PF synapses, the direction and size of LTP or LTD are determined by the combination of multiple simultaneous LTP and LTD mechanisms (O’Connor *et al.*, 2005; Bender *et al.*, 2006; Tzounopoulos *et al.*, 2007; Shen *et al.*, 2008; Zhao & Tzounopoulos, 2011). In DCN PF synapses, LTP and LTD are influenced by the coactivation of pre-and postsynaptic signaling mechanisms including NMDARs, mGluRs, muscarinic acetylcholine receptors, and endocannabinoid signaling (Fujino & Oertel, 2003; Tzounopoulos *et al.*, 2007; Zhao & Tzounopoulos, 2011). Therefore, bidirectional zinc plasticity likely acts together with these other known mechanisms to shape the size and direction of synaptic plasticity.

Several of our results are consistent with this notion. As shown in Figure 2, blockade of NMDARs did not block either HFS-induced LTP or Z-LTD. This indicates that NMDAR-independent LTP was induced, suggesting that Z-LTD contributes to NMDAR-independent LTP (Figure 2 A). G1 mGluR antagonists blocked HFS-induced Z-LTD (Figure 2 B). Therefore, the induced LTP under these conditions is NMDAR-, G1 mGluR-, and Z-LTD-independent. As shown in Figure 3 A, LFS induced Z-LTP; however, LFS did not induce LTD. This suggests that LFS also induced an LTP that counterbalances the LTD effect of Z-LTP. This is consistent with previous studies showing that LTP and LTD mechanisms occur simultaneously in DCN PF synapses (Tzounopoulos *et al.*, 2007). G1 mGluR antagonists blocked Z-LTP, but LTD was induced under these conditions (Figure 3 B), suggesting that this LTD is NMDAR-, G1 mGluR-, and Z-LTP-independent. Taken together, all these results are consistent with previous studies and further support the notion that LTP and LTD are the result of coactivation of different signaling pathways of long-term plasticity in the DCN (Fujino & Oertel, 2003; Tzounopoulos *et al.*, 2007; Zhao & Tzounopoulos, 2011). Nevertheless, our results add Z-LTP and Z-LTD as new mechanisms of LTD and LTP at zinc-containing glutamatergic synapses.

In DCN PFs, mGluR activation contributes to both HFS-induced LTP and LFS-induced LTD (Fujino & Oertel, 2003). Our findings on HFS-induced LTP in CWCs are consistent with Fujino & Oertel, 2003 (Fujino & Oertel, 2003), demonstrating mGluR-and NMDAR-independent LTP (Figure 2 B). However, Fujino & Oertel showed LFS-induced NMDAR-dependent LTD, whereas here we observed LFS-induced NMDAR-independent LTD (Figure 3 B). This discrepancy could be explained by the use of different LFS induction protocols (5 Hz for 3 min here, vs. 1 Hz for 5 min. paired with postsynaptic depolarization) and extracellular solutions (4/4 mM Ca^2+^/Mg^2+^ here, vs. 2.4/1.3 mM Ca^2+^/Mg^2+^).

In addition to DCN PF synapses, Z-LTP and Z-LTD may contribute to LTD and LTP in other synaptic zinc-containing brain areas which express G1 mGluR-dependent LTD and LTP, such as the hippocampus, amygdala, and striatum (Oliet *et al.*, 1997; Huber *et al.*, 2000; Gubellini *et al.*, 2003; Topolnik *et al.*, 2006; Wu *et al.*, 2008; Anwyl, 2009; Luscher & Huber, 2010; Chen *et al.*, 2017; McAllister & Dyck, 2017). In the hippocampus, LFS induces G1 mGluR-mediated LTD, whereas HFS induces LTP (Oliet *et al.*, 1997; Huber *et al.*, 2000; Topolnik *et al.*, 2006; Wu *et al.*, 2008; Anwyl, 2009). Therefore, LFS-induced Z-LTP would likely further enhance the effects of G1 mGluR-LTD, by increasing zinc inhibition of AMPARs; whereas HFS-induced Z-LTD would further enhance the effects of LTP, by reducing zinc inhibition of AMPARs. Thus, synaptic zinc plasticity likely serves as a positive feedback mechanism to enhance the effects of G1 mGluR-dependent LTP or LTD on glutamatergic synaptic transmission.

### Implications of Z-LTP and Z-LTD for metaplasticity

Our results reveal that the induction of Z-LTP and Z-LTD is NMDAR-independent. However, zinc inhibits NMDARs and thus modulates the induction of NMDAR-dependent LTP and LTD in the hippocampus (Izumi *et al.*, 2006; Takeda *et al.*, 2009; Vergnano *et al.*, 2014). As such, long-term synaptic zinc plasticity may contribute to ‘metaplasticity’, the modulation of subsequent LTP and LTD (Abraham & Tate, 1997). Z-LTD, by reducing the inhibitory effect of zinc on NMDARs, may promote subsequent NMDAR-dependent LTP and decrease subsequent NMDAR-dependent LTD. Conversely, Z-LTP, by enhancing the inhibitory effect of zinc on NMDARs, may promote subsequent NMDAR-LTD over NMDAR-LTP. Therefore, zinc plasticity likely serves as a positive feedback mechanism for NMDAR-dependent metaplasticity.

Synaptic zinc contributes to mossy fiber presynaptic LTP in response to HFS, via activation of TrkB receptors (Huang *et al.*, 2008; Pan *et al.*, 2011). Therefore, if HFS induces Z-LTD in mossy fiber synapses, it would act as a negative feedback signal for subsequent LTP induction. Taken together, we propose that the role of Z-LTD and Z-LTP in LTP and LTD depends on the specific mechanisms underlying LTP and LTD, but overall, Z-LTD and Z-LTP likely act as positive feedback signals in G1 mGluR-dependent synaptic plasticity and NMDAR-dependent metaplasticity.

### Clinical and translational implications of zinc plasticity

In the context of zinc plasticity as a positive feedback signal for NMDAR-dependent metaplasticity, it is interesting that exposure to loud sound – known to induce tinnitus – causes Z-LTD in the DCN. Although not tested here, it is possible that Z-LTD could potentially lead to runaway excitation due to enhanced LTP and decreased LTD, and thus to pathological DCN hyperactivity associated with tinnitus (Tzounopoulos, 2008). Noise-induced pathological hyperexcitability through LTP/LTD-like mechanisms in the DCN PF synapses has been hypothesized and recently implicated in tinnitus treatment (Tzounopoulos, 2008; Marks *et al.*, 2018), therefore suggesting that noise-induced reductions in synaptic zinc might contribute to tinnitus.

## Funding and Acknowledgements

This work was supported by NIH grants F31-DC015924 (to N.W.V.) and R01-DC007905 (to T.T.). We thank Drs. Stephen Lippard and Jacob Goldberg from Massachusetts Institute of Technology for generously providing DA-ZP1. We thank Patrick Cody and Dr. Ross Williamson for technical help with data analysis. We thank Dr. Elias Aizenman for helpful discussions. The authors declare no competing financial interests.

